# Microbiome evolution plays a secondary role in host rapid adaptation

**DOI:** 10.1101/2025.06.27.661976

**Authors:** René S. Shahmohamadloo, Amir R. Gabidulin, Ellie R. Andrews, Seth M. Rudman

## Abstract

Understanding how populations adapt to environmental change is a central goal in evolutionary biology. Microbiomes have been proposed as a source of heritable variation that is central to rapid adaptation in hosts, yet empirical evidence supporting this remains limited, particularly in naturalistic settings. We combined a field evolution experiment in *Drosophila melanogaster* exposed to an insecticide with microbiome manipulations to disentangle the contributions of host standing genetic variation and microbiome evolution to adaptation. Within three generations, independent populations rapidly and repeatedly evolved increased survivorship, a defining feature of resistance evolution. Adaptive changes in sub-lethal traits such as reproductive output, stress tolerance, and body size occurred with a delayed response following the evolution of resistance. Core microbiome taxa declined following insecticide exposure, and resistant populations evolved to house lower microbial abundances. Axenic rearing and microbiome transplant experiments demonstrated that adaptation via host standing genetic variation was the mechanism for resistance evolution. Microbiome evolution played a secondary and cryptic role in host adaptation by masking slowed development rates that evolved in resistant populations. Together, these results reinforce the primacy of adaptation occurring through selection on host standing genetic variation while also demonstrating the contributions of microbiome evolution in host adaptation.

**Significance:** Identifying the mechanisms that allow organisms to adapt to environmental stress is a foundational goal in biology. Using field experimental evolution and microbiome manipulations in *Drosophila melanogaster*, we directly tested the relative contributions of host genomic evolution and microbiome evolution to adaptation. We found that adaptation to environmental stress occurred rapidly and repeatedly, driven primarily by selection on host standing genetic variation, with microbiome evolution acting as a secondary contributor. These findings reinforce the importance of host genetic variation in rapid adaptation and demonstrate that microbiome evolution can contribute to host evolutionary trajectories in a cryptic manner.

## Introduction

Understanding how populations adapt to environmental change remains a central challenge in evolutionary biology (1–3) with critical implications for the preservation of biodiversity in the Anthropocene (4, 5). Adaptation depends on both the strength of selection imposed by environmental stressors and the availability of additive genetic variation in traits under selection (6, 7). Studies of rapid evolution in eukaryotes have identified standing genetic variation as the substrate on which selection acts to drive adaptation (7–9). Experimental evolution studies and theoretical models have demonstrated that adaptation from standing genetic variation can enable populations to persist in changing environments (1, 6). Additive genetic variation may derive from multiple sources—including the host genomic sequence (10), transgenerational epigenetic inheritance (11), and host-microbe interactions (12)—yet the relative contributions of these mechanisms to adaptation remains poorly understood.

Microbiomes have received particular attention for their potential to contribute to host adaptation, not only through effects on host physiology but also by influencing heritable trait variation in some taxa when transmitted across generations (12–15). The *holobiont* framework proposes that hosts and their associated microbes can act as integrated units of selection (16–19). However, empirical support for microbiome-driven adaptation comes primarily from systems of obligate symbiosis (20–24). Many organismal microbiomes are primarily colonized from environmental exposure, and the degree to which environmentally acquired microbiomes contribute to host evolution remains unclear (25, 26).

Microbiome evolution—defined as heritable change in microbial composition across host generations (27)—is central to the *holobiont* theory of evolution. For microbiome evolution to influence host phenotypes, microbial taxa generally need to be stably transmitted either vertically or via host-mediated horizontal acquisition, and affect traits that contribute to host fitness (13, 28). Many studies have shown that microbiome composition shifts in response to environmental stress (24, 29, 30), and some have further demonstrated that these shifts can influence host performance (31–34). Few studies extend to testing whether shifts in microbiome composition are heritable (31), and examinations of the contribution of microbiome evolution on host adaptation are rare. Moreover, studies that reported evidence of microbiome evolution often restricted or eliminated genetic variation in the host population (31, 32, 34, 35). This facilitates isolation of the effects of the microbiome on host evolution but precludes assessing microbiome contributions relative to other sources of heritable variation shaping host evolution. Such experimental systems do not reflect the complexity of many natural eukaryotic populations, where variation exists in both host genotype and microbiome composition. As a result, they may overestimate the role of microbiomes in host adaptation and in population responses such as evolutionary rescue (36), or fail to capture how microbiome evolution unfolds under more ecologically realistic conditions (37).

Experimental evolution in natural systems offers a potentially powerful approach for testing whether microbiome evolution contributes to host adaptation (14). Field evolution experiments have been critical to our understanding of the pace of evolution and the action of natural selection (38–41), including studies tracking temporal evolution of *Drosophila melanogaster* (42–44). Combining a replicated field evolution experiment that identifies adaptation through parallel phenotypic change across independent populations with microbiome manipulations, including removal and transplant experiments, provides a powerful framework to test whether adaptation occurs through evolution of host standing genetic, microbiomes, or both.

To empirically test the contribution of microbiome evolution to host adaptation, we introduced outbred *D. melanogaster* populations into 2m × 2m × 2m outdoor enclosures where they evolved on a control (no insecticide) or toxic (insecticide-treated) diet (Figure 1). Over nine generations, we tracked population demography, host phenotypic evolution, and microbiome composition. We examined both the effects of insecticide exposure on microbiome composition and the impact of resistance evolution on microbiome evolution. Finding evidence for both effects of exposure and evolutionary divergence in microbiome composition, we repeated this experiment but with the addition of microbiome removal and transplant experiments to disentangle the contribution of the microbiome to host resistance evolution. This design allowed us to answer four core questions: 1) Do *D. melanogaster* populations evolve resistance to insecticide exposure, and if so, what was the pace, phenotypic basis, and predictability of this evolutionary response? 2) What was the magnitude and parallelism of microbiome shifts associated with insecticide exposure? 3) Did microbiomes evolve as host populations evolved resistance? 4) Did microbiome evolution contribute to host phenotypic adaptation? *D. melanogaster* is well-suited for this work due to its short generation time, genetic diversity, and well-characterized microbiome with known effects on host fitness traits including development rate and stress tolerance (12, 45–48). Among selective agents, insecticides are particularly compelling because they exert strong selection on hosts and are known to alter microbial community composition. Insects exposed to agrochemicals frequently exhibit shifts in their microbiomes, with documented consequences for host physiology and stress responses. Moreover, microbiome composition has been implicated as a potential mechanism of insecticide resistance evolution in insects (), and resistance evolution (32, 49–52). To disentangle host and microbiome contributions to adaptation, we used repeated common garden rearing of 10 field experimental populations to track adaptation. We conducted within generation exposure to the insecticide to test for the direct effects of insecticide exposure on the microbiome. We used common garden rearing and subsequent microbiome sequencing to test for divergent microbiome evolution between insecticide-exposed and control populations. We compared the magnitude of resistance evolution, both in lethal (e.g., survivorship) and sub-lethal (e.g., fecundity, starvation tolerance, adult weight) traits, with and without the microbiome present (e.g., axenic rearing) and used gnotobiotic transplant experiments that introduced gut microbiota from resistant and control populations into naïve hosts to test whether microbiome evolution contributed causally to host adaptation.

**Figure 1.**
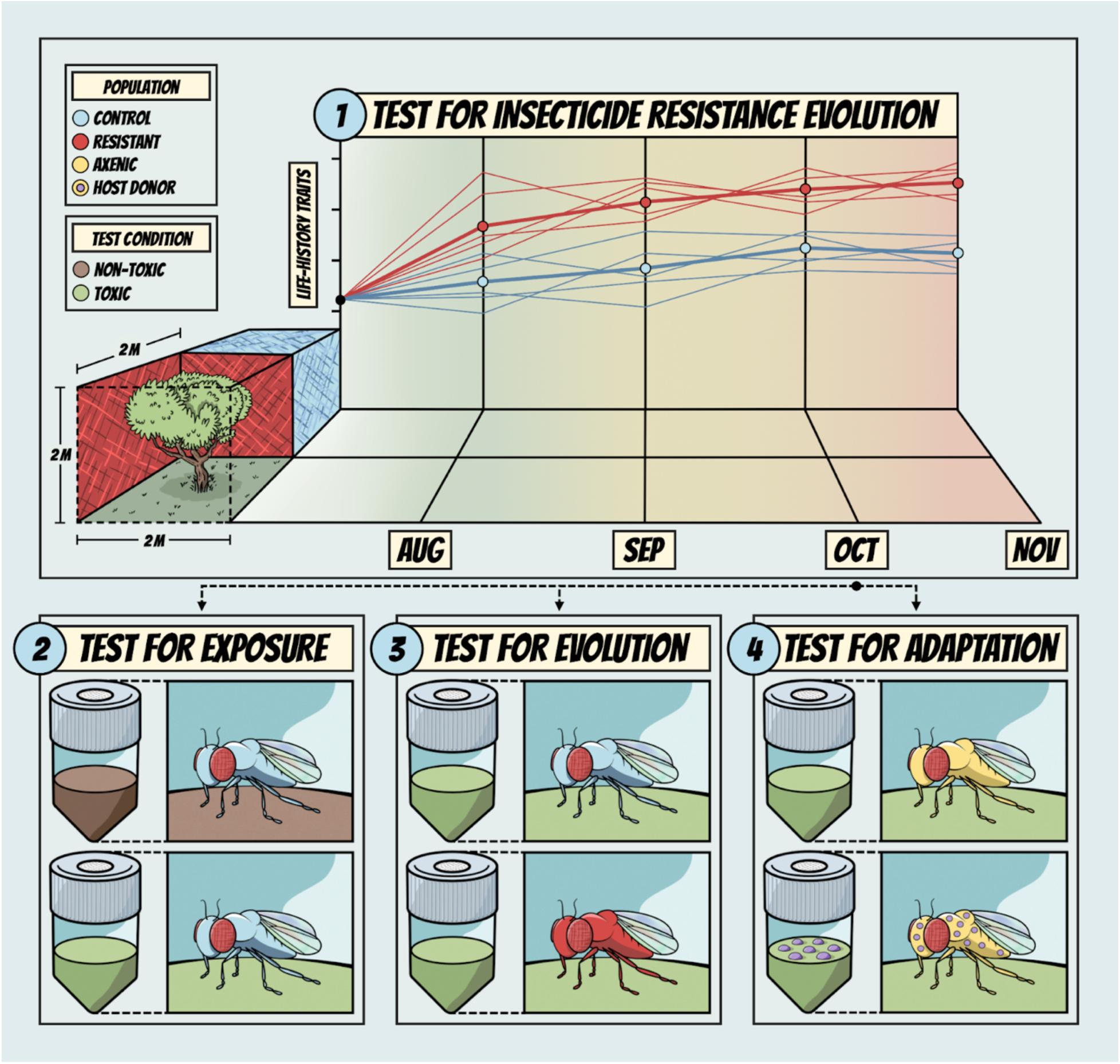
Replicate outdoor mesocosms containing *Drosophila melanogaster* populations were exposed to non-toxic (no insecticide) and toxic (insecticide) conditions from mid-summer to late-fall to: (1) Test whether *D. melanogaster* populations evolve resistance to insecticide exposure, including both lethal and sub-lethal traits. In October, eggs from each population were collected and reared in common garden conditions for two generations to: (2) Test for the effects of insecticide exposure on microbiome composition; (3) Test for microbiome evolution associated with host resistance evolution by rearing descendants from each field cage in a common garden environment and examining differences in microbiome composition; and (4) Test for the contribution of microbiome evolution to host adaptation through two complementary approaches: (top panel) axenic rearing (microbiome removal) compared to xenic rearing to assess microbiomes contribution to resistance evolution; and (bottom panel) microbiome transplant experiments, where microbiota from control and resistant donor populations were introduced into germ-free hosts.

## Results

### Phenotypic evolution

To assess the effects of insecticide exposure on fitness-related traits, we measured survivorship, fecundity, starvation tolerance, adult weight, and development rate—key life-history traits that influence population persistence under environmental stress (Figure 2). All five traits evolved rapidly and in parallel over the course of the experiment in both control and resistant populations when assayed on insecticide media (survivorship, F_4,32_ = 29.48, P < 0.0001; fecundity, F_4,32_ = 11.90, P < 0.0001; starvation tolerance, F_4,40_ = 10.93, P < 0.0001; adult weight, F_4,34_ = 16.69, P < 0.0001; development rate, F_4,32_ = 7.09, P = 0.0003).

**Figure 2.**
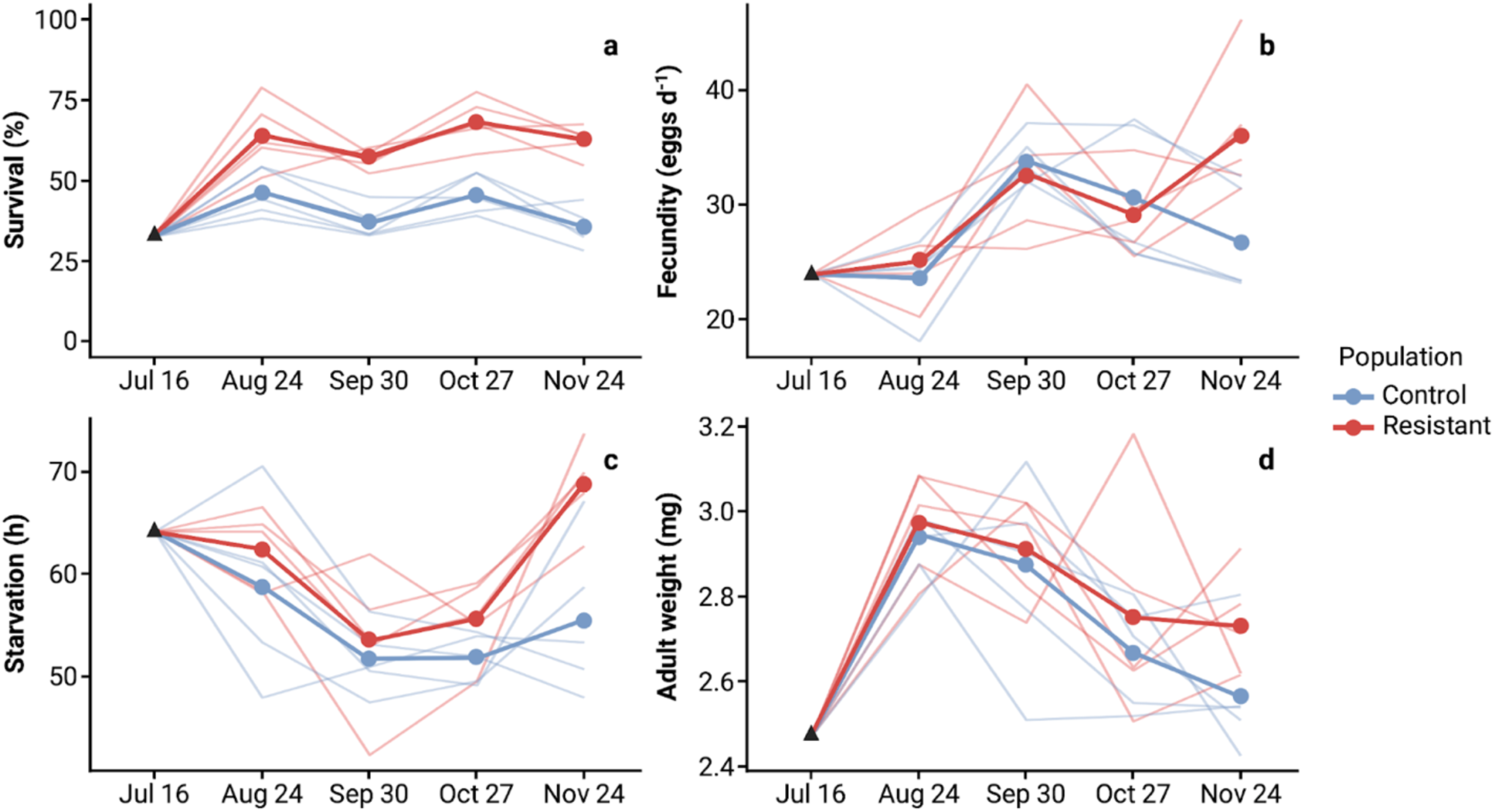
Phenotypic evolution of life-history traits in populations exposed (‘resistant’) and unexposed (‘control’) to the insecticide. Phenotypic trajectories were measured at each time point following two generations of common garden rearing, with all assays conducted on insects reared on toxic (insecticide) media to assess resistance. (A) Mean survival, measured as the percentage of eggs (30 per vial) that survived to adulthood. (B) Mean fecundity, measured as the number of eggs produced per female per day. (C) Mean starvation tolerance, measured as the time to death by starvation. (D) Mean adult weight, measured as the average dry weight of female flies. Black triangles (▴) represent the mean phenotype of the founding population (outdoor cages initiated July 16), measured under toxic assay conditions. Thin colored lines represent the mean phenotypic trajectory of each individual population, and thick colored lines show the treatment means (averaged across cages). Mean development rate, calculated as the fraction of development time completed per day [1/(total hours/24)], can be found in the Supporting Information (Figure S1).

Insecticide exposure led to parallel evolutionary divergence between insecticide and control populations indicative of adaptation in three of the five phenotypes, as indicated by a significant Time × Population interaction for survivorship (F_4,32_ = 8.36, P < 0.0001), fecundity (F_4,32_ = 3.36, P = 0.021), and starvation tolerance (F_4,40_ = 2.66, P = 0.046). The effect of insecticide exposure on survival was particularly strong, with resistant populations exhibiting 39% higher survivorship than controls after approximately three generations of exposure (Figure 2). This divergence in survivorship was driven by faster evolutionary change in resistant populations, which evolved at 0.89 haldanes during the initial interval (*T*_0_ → *T*_1_; Supporting Information, Table S1). In contrast, divergence in fecundity and starvation tolerance emerged more gradually, with resistant populations ultimately exhibiting 34% and 18% higher values than controls by the end of the experiment (Figure 2). Notably, the largest rates of evolutionary change in sub-lethal traits occurred in the latter half of the experiment, with resistant populations showing elevated rates in fecundity (*T*_3_ → *T*_4_: 0.130 haldanes) and starvation tolerance (*T*_3_ → *T*_4_: 0.390 haldanes), compared to controls (fecundity = −0.075 haldanes; starvation tolerance = 0.116 haldanes; Supporting Information, Table S1), consistent with accelerated divergence over time.

### Microbiome shifts and evolution under insecticide stress

To assess the effects of insecticide exposure, we compared microbial diversity and composition in the founder population reared on non-toxic (no insecticide) versus toxic (insecticide) media (Figure 3a). Within-generation exposure to the insecticide led to detectable shifts in microbiome composition (Weighted UniFrac: F_1,18_ = 4.27, P = 0.026; Bray-Curtis: F_1,18_ = 1.37, P = 0.20). We quantified the direction of microbiome shifts using angular difference metrics (53) and found that changes in composition across independent populations were more parallel than expected by chance with angles of 82.78° for Weighted UniFrac (t_17_ = −1.81, P = 0.087) and 82.35° for Bray-Curtis (t_17_ = −3.19, P = 0.005). These results indicate some parallelism in the way microbiome communities shift in response to insecticide exposure. Insecticide exposure also reduced the abundance of the two dominant bacterial families: *Acetobacteraceae* by 25% (Z_1,18_ = −68.77, P < 0.001), and *Lactobacillaceae* decreased by 18% (Z_1,18_ = −5.9, P < 0.001).

**Figure 3.**
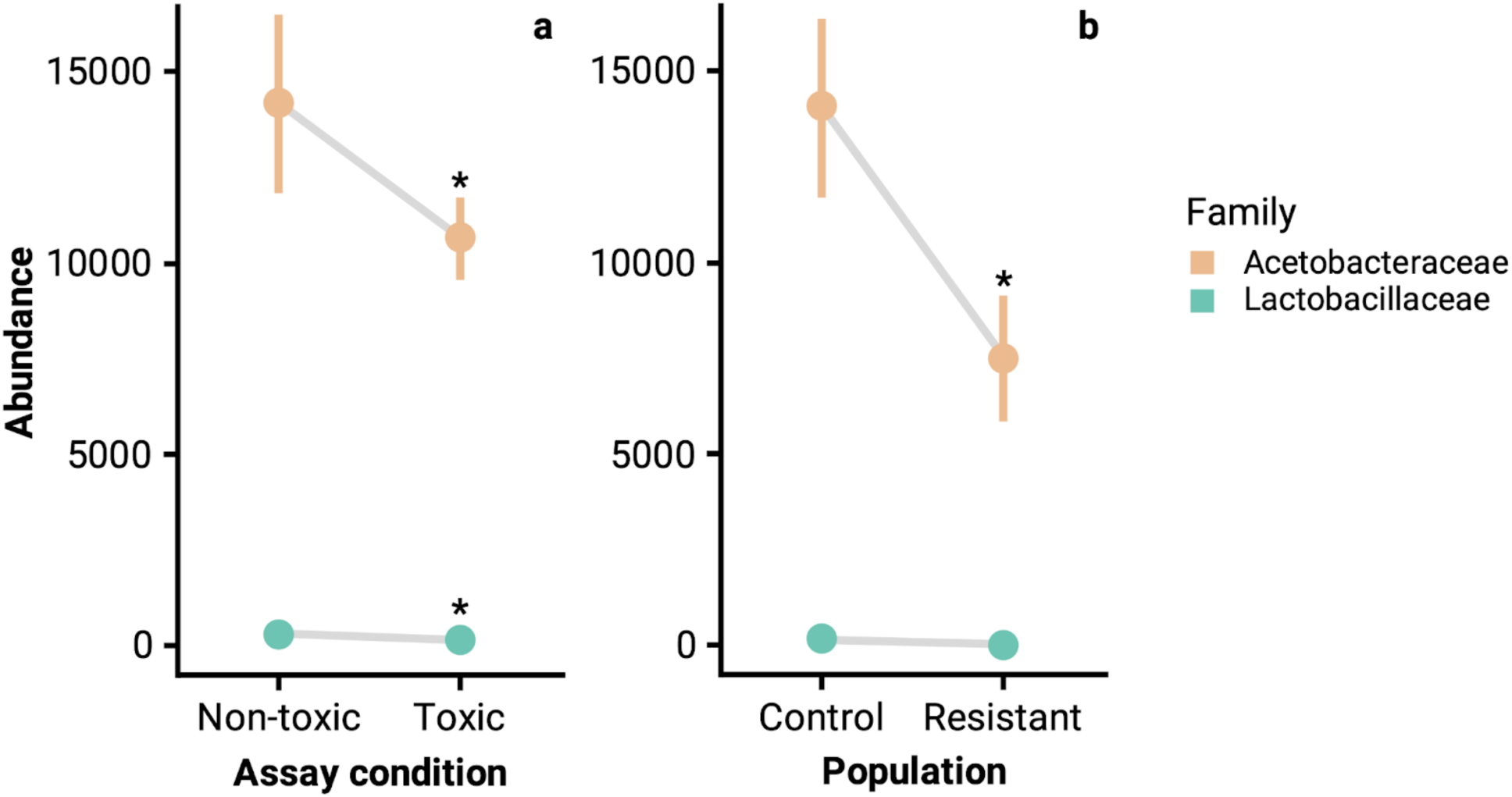
Effects of insecticides on the microbiome measured both through direct exposure and evolutionary divergence associated with the evolution of resistance. Mean abundance (±1 SE) of the two most dominant microbial families, *Acetobacteraceae* and *Lactobacillaceae*, in *Drosophila melanogaster* populations exposed to insecticide stress. (A) To evaluate the microbiome’s response to insecticide exposure, microbiomes of the founding population were assayed after within-generation exposure to non-toxic (no insecticide) and toxic (insecticide) conditions, after two generations of common garden rearing. (B) To evaluate microbiome evolution, microbiomes of control and resistant populations were compared after two generations of common garden rearing on toxic media. Significant differences in abundance between populations are designated by an asterisk (*). The full list of microbial families can be found in the Supporting Information Table S2.

To assess microbiome evolution, we tested for differences in microbial diversity and composition between resistant and control populations following two generations of common garden rearing on toxic media (Figure 3b). Beta diversity analyses revealed no significant differences between resistant and control populations in microbiome composition (Weighted UniFrac: F_1,16_ = 2.156, P = 0.126; Bray-Curtis: F_1,16_ = 2.536, P = 0.059). However, we did observe parallelism in shifts of microbial composition associated with resistance evolution Weighted UniFrac mean angle was 73.65° (*t*_15_ = −4.74, P < 0.001) and 84.34° for Bray Curtis (*t*_15_ = −1.71, P = 0.108) (Supporting Information, Figure S2). Moreover, resistant populations exhibited large reductions in the abundance of the two dominant microbial families: *Lactobacillaceae* abundance was 82% lower (Z_1,16_ = −27.4, P < 0.001), and *Acetobacteraceae* was 46% lower (Z_1,16_ = −126.8, P < 0.001), suggesting that adaptation to toxic conditions was associated with microbiome evolution. In particular, resistant populations did evolve to host lower abundances of *Lactobacillus*, a well-studied taxon known to affect host physiology, particularly under toxic conditions (non-toxic: *t*_15,16_ = −1.648, P = 0.12; toxic: *t*_37,38_ = −2.585, P = 0.014).

### Contribution of the microbiome to host phenotypic adaptation

To assess whether the microbiome contributed to resistance evolution, we compared phenotypes of flies reared in axenic (microbiota-removed) versus xenic (microbiota-present via natural recolonization) conditions. Resistance evolution, as measured by survival differences between resistant and control populations, was not significantly impacted by microbiome treatment; resistant populations maintained higher survival under both axenic and xenic conditions (F_1,8_ = 1.10, P = 0.33; 95% CI = −6.79, 2.55; Figure 4a). Similarly, adult weight did not significantly differ across microbiome treatments (F_1,8_ = 0.03, P = 0.87; 95% CI = −0.41, 0.36; Figure 4b), suggesting that the increased body size observed in resistant populations was an evolved host response to insecticide exposure rather than a microbiome-mediated effect. Development rate, however, was more strongly reduced in resistant populations when reared axenic (F_1,8_ = 56.71, P < 0.001; 95% CI = 0.00624, 0.0118; Figure 4c). This shift corresponds to a delay of approximately 21 hours in development time, a potentially biologically meaningful impact of the microbiome on developmental timing under insecticide stress.

**Figure 4.**
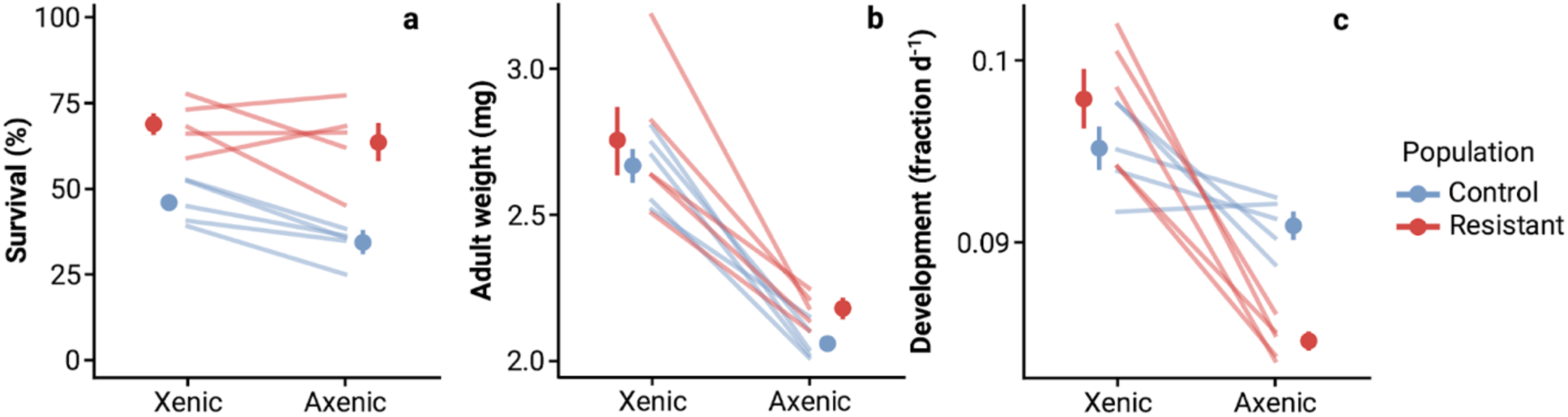
Testing whether microbiome evolution contributed to host adaptation. Panels A-C show trait values for resistant and control populations reared on a toxic diet under either xenic (microbiome present) or axenic (microbiome removed) conditions. Points represent cage-level means, and lines connect cages that were paired based on spatial proximity in the outdoor experimental system. Bold points and error bars indicate the mean ± SE across cages. (A) Mean survival, measured as the percentage of eggs (30 per vial) that survived to adulthood. (B) Mean adult weight, measured as the average dry weight of female flies. (C) Mean development rate, calculated as the fraction of development time completed per day [1/(total hours/24)]. Axenic flies were generated by sterilization and reared in bioreactor tubes as gnotobiotic flies.

As an additional test of whether microbiome evolution contributed to host adaptation, we conducted a transplant experiment to determine whether any divergent microbiome evolution between resistant and control populations could recover the observed patterns of host resistance evolution. Individuals from each resistant and control population were reared for two generations in common garden conditions before being used as microbiome donors onto pools of axenic eggs taken from the founding population.

The transplant experiment revealed that microbiome composition was not significantly impacted by donor population (control vs. resistant; F_1,19_ = 1.9083, P = 0.124; Figure 5a). However, the donor population clearly impacted the abundance of *Acetobacteraceae* (non-toxic: *z*_8,9_ = 121.2, P < 0.001; toxic: *z*_8,9_ = 71.16, P < 0.001) and *Lactobacillaceae* (non-toxic: *t*_8,9_ = 6.297, P < 0.001; toxic: *t*_8,9_ = −3.105, P = 0.0019) on non-toxic and toxic diets. However, these difference in abundance did not translate to measurable phenotypic effects for the host; whether donor microbiomes came from control or resistant populations had no detectable effect on host survival (*t* = −0.264, P = 0.994; 95% CI = −2.02, 1.56; Figure 5b), development rate (*t* = −1.010, P = 0.7433; 95% CI = −1.073. 0.468; Figure 5c), or adult weight (*t* = −1.335, P = 0.5427; 95% CI = −0.278, 0.053; Figure 5d).

**Figure 5.**
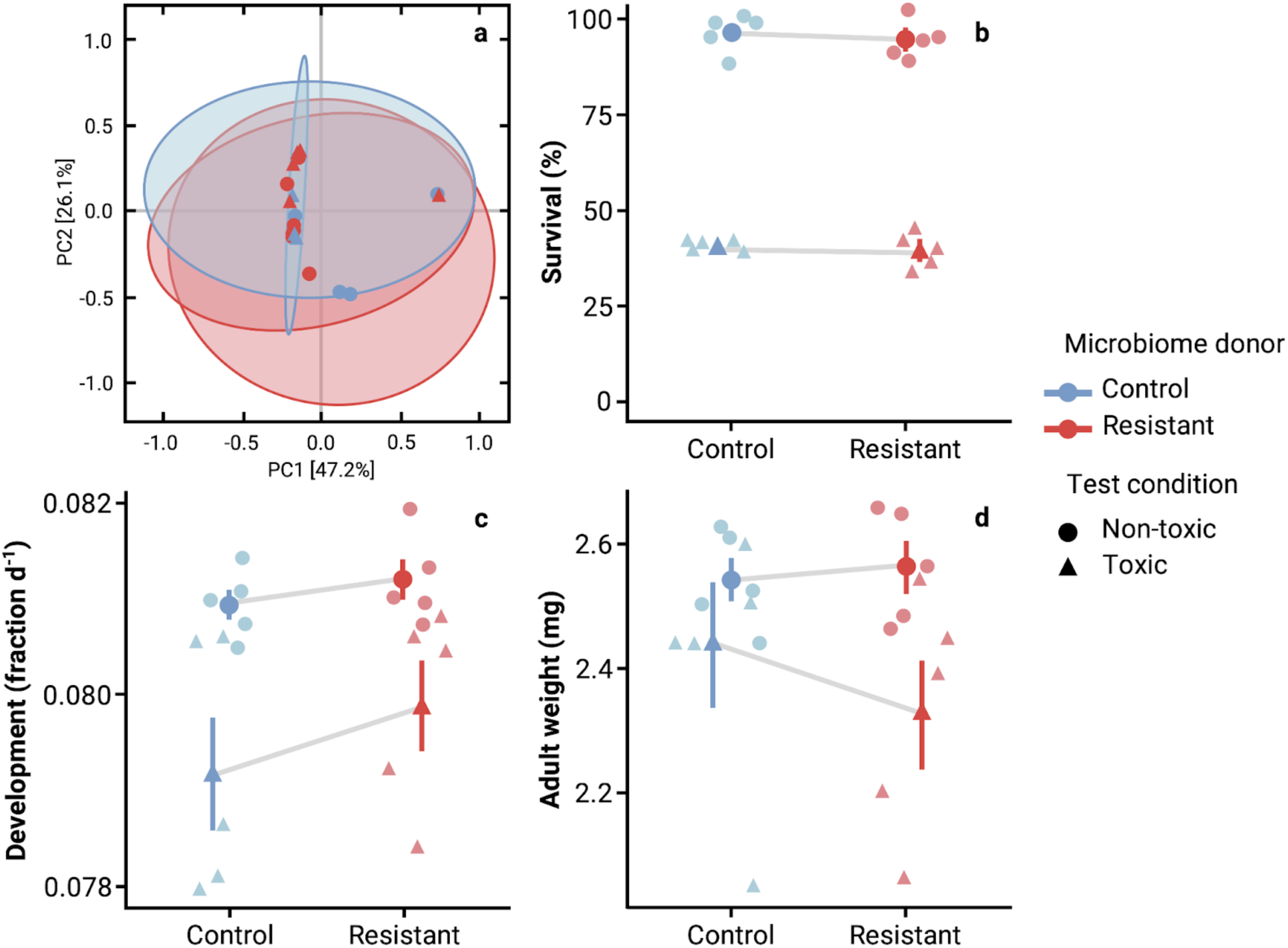
Testing for the contribution of microbiome donors to host adaptation. Panels (A-D) show the results of microbiome transplants in which microbiomes from each of the 10 outdoor populations (five resistant, 5 control) were transplanted into subsets of the ‘founder’ population. Founder flies received gut microbiomes of either a pooled homogenate from common garden reared control or resistant populations. These recipient flies were measured to determine whether microbiome transplants could recover patterns of phenotypic adaptation. (A) Principal coordinate analysis (PCoA) of Bray-Curtis distances showing microbiome composition of recipient flies after transplants from control (blue) or resistant (red) donor populations; microbiome composition did not differ significantly by donor type. (B) Mean survival as the proportion of eggs (30 eggs per vial) that survived to adulthood; no significant difference by microbiome donor. (C) Mean development rate as the time at which flies eclosed (30 eggs per vial); no significant difference by microbiome donor. (D) Adult weight as the average dry weight of female flies. All flies were sterilized and reared in bioreactor tubes as gnotobiotic flies to complete all assays (B-D).

## Discussion

### Divergent timescales of trait evolution under environmental stress

The rapid evolution we observed in the lethal trait of survivorship (Figure 2a) reflects strong selection acting directly on viability, where resistance to a lethal dose at which 50% of a population experiences mortality (LD50; (54, 55)) evolves quickly under intense selection pressure (56–60). By comparison, the gradual evolution in the sub-lethal resistance traits of fecundity (Figure 2b) and starvation tolerance (Figure 2c) when exposed to insecticides suggests that additional physiological adaptations unfold over longer timescales following viability selection, potentially due to trade-offs or tolerance mechanisms that only become apparent—or become subject to selection—after resistance is established (58, 61, 62). This pattern aligns with perspectives from evolutionary toxicology (59) and evolutionary genetics (1), where acute responses to environmental challenges are often detectable early, but longer-term adaptations involving physiological trade-offs unfold over extended timescales. Clarifying the generality of these divergent evolutionary timescales could provide new insights into the dynamics of resistance evolution and the long-term efficacy of insecticide resistance management strategies.

### Cryptic and secondary role of microbiome evolution in host adaptation

Insecticide exposure rapidly reduced the relative abundances of *Acetobacteraceae* and *Lactobacillaceae* (Figure 3a), a pattern consistent with prior evidence in insects that pesticides can disrupt gut microbiota and suppress core microbial taxa (52, 63, 64). We observed a similar pattern associated with microbiome evolution; *Acetobacteraceae* abundance was reduced in resistant populations (Figure 3b). *Acetobacteraceae* and *Lactobacillaceae* have previously been linked to spatial adaptation in *Drosophila* and shown to influence host life-history trade-offs (47, 48). Specifically, acetic acid bacteria are associated with reduced stress tolerance and increased reproductive output, while lactic acid bacteria can drive opposite phenotypic shifts (48). Despite limited changes in overall beta diversity, the consistent and parallel reductions in *Acetobacteraceae* and *Lactobacillaceae* suggest deterministic, selection-driven shifts in the microbiome during resistance evolution. These findings suggest that microbiome shifts observed in wild insect populations following insecticide exposure may, in part, reflect underlying host resistance evolution, not solely the direct effects of chemical exposure.

Despite clear effects of resistance evolution on the abundance of key microbiome taxa, microbiome transplant experiments showed that these evolved differences in microbiome communities provided no fitness benefit to naïve hosts and had no direct contribution to lethal resistance (Figure 5). However, axenic rearing revealed that the microbiome buffered a fitness cost in development rate: resistant populations developed significantly slower in the absence of their microbiome (∼21 hours). Microbiome presence restored developmental timing to rates similar to those observed in control populations (Figure 4c). Evolutionary divergence in development rate among insects is commonly associated with adaptation to varying environmental conditions, including both spatial and temporal variation (65, 66). While the microbiome did not drive adaptation directly, its ability to buffer a developmental cost of resistance suggests a facilitative role in masking a potential phenotypic trade-off between resistance and development. This masking effect of the microbiome could help maintain beneficial resistance alleles under stress. Even when adaptation is primarily driven by host genetic changes, the microbiome can influence evolutionary outcomes by modulating the expression of evolved traits (13, 25). This illustrates a cryptic eco-evolutionary dynamic (67), in which microbial interactions influence how costs of adaptation are expressed.

### Host genetic variation drives adaptation

Our findings underscore a growing distinction between microbiomes as ecological agents that shape host phenotypes (13, 48, 52, 68) and as a component of heritable variation that contributes to host adaptation (17, 25, 69–72). This outcome aligns with theoretical models showing that for microbiomes to evolve as extensions of host genetic architecture, their community dynamics must be governed more by vertical transmission and within-host selection rather than by environmental filtering and stochastic assembly, and they must generate repeatable, fitness-relevant effects on host phenotypes (69–71, 73, 74). While we observed some deterministic shifts in microbial composition during resistance evolution, the full set of conditions required for microbiomes to act as primary agents of host adaptation were not met in this system. In settings where microbiota are environmentally acquired, microbiomes may more often function as plastic responders to environmental change and capable of influencing evolutionary outcomes even when not serving as a primary source of heritable variation on which selection can act (14, 75).

This distinction has important implications. If microbiomes are to be included in evolutionary models of host adaptation, their transmission mode, stability, and trait-specific effects must be empirically established. Without such criteria, microbiome evolution risks being misinterpreted as evidence of host adaptive evolution. Our findings suggest that while microbiomes can buffer or modulate trait expression in important ways, their role as the substrate on which selection acts to drive host adaptation—at least under short timescales and strong directional selection—may be modest in horizontally acquired systems such as *Drosophila*.

## Materials and Methods

### Founding population

To establish the founding *D. melanogaster* population, we crossed 100 isofemale lines from the Drosophila Genetic Reference Panel (DGRP) (76), following previous methodologies (44, 47). Ten males and 10 females from each line were combined in a breeding cage, and six generations of random mating were conducted under density-controlled laboratory conditions (25°C, 12L:12D photoperiod, and fed a modified ‘Bloomington Drosophila Stock Center Recipe’ media) to facilitate recombination before founding the outdoor mesocosm experiments.

### Outdoor experimental design

Replicated outdoor mesocosm experiments were conducted in 2022 and 2023 using 2m × 2m × 2m mesh enclosures constructed of fine mesh built around metal frames (BioQuip PO 1406C). Each mesocosm contained a non-fruiting dwarf apple tree and vegetative ground cover to provide shading and microbial communities (44, 47).

Flies were introduced into mesocosms on July 17, 2022, and July 16, 2023, respectively. For both experiments, 2,500 F6 flies of a single age cohort were seeded into each mesocosm. The insecticide chosen for our experiments was spinosad, a widely used organic insecticide known for its mechanisms of action in pests and documented resistance impacts in Drosophilid flies (77, 78). Spinosad dosages were determined using dose-response curves for egg-to-adult survivorship (Supporting Information, Figure S3). In 2022, populations were split among three treatments: a non-toxic control and two insecticide treatments at concentrations of 0.0188 µg L_-1_ and 0.0375 µg L_-1_ spinosad, with five replicates per treatment, totalling 15 mesocosms. In 2023, we refined the experimental design to include one control and a single insecticide treatment of 0.0375 µg L_-1_ spinosad, with five replicates per treatment, totalling 10 mesocosms. Food media was provided in 400 mL increments in 900 cm³ aluminum loaf pans every two days as the sole food source and egg-laying substrate for each mesocosm. Food was sheltered from direct rain and sunlight. After two days, each pan was covered with a mesh lid to prevent further egg laying, and monitored daily until fly eclosion when they were released (43, 44).

### Phenotypic assays

To assess the evolutionary response to insecticide exposure, ∼750 eggs were collected from each replicate mesocosm at regular intervals (August, September, October, November) and reared under common garden conditions (25°C, 12L:12D photoperiod) for two generations (F0→F2). This ensured that observed phenotypic differences were genetic rather than environmental. The following phenotypes were assayed per replicate mesocosm: 1) Survivorship to adulthood, measured as the proportion of eggs (30 eggs per vial) that survived to adulthood; 2) Development rate, measured as the time from egg to pupation (i.e., 1/time of development from egg to adult); 3) Fecundity, measured as the total eggs laid by five females over three days; 4) Starvation tolerance, measured as the time to starvation for three replicate vials containing 10 males each on agar-only media; 5) Adult weight, measured the average weight of three pools of five females, dried at 55°C for 24 h. Population sizes were estimated weekly by photographing four, 0.3 m_3_ transects on the ceiling of each mesocosm near sunset when flies aggregate on the walls, followed by using a custom trained machine learning algorithm to count the flies in each photo (79).

### 2 × 2 experimental design and transplant

We conducted a 2 × 2 factorial experiment in 2023 using eggs collected on October 27, 2023, and reared in an indoor common garden for three generations prior to use. Eggs from control and resistant populations were sterilized by bleaching (two 150-second rinses) in 0.6% sodium hypochlorite and washed three times in sterile distilled water before being transferred to sterile media in bioreactor tubes (30 eggs per tube) to produce gnotobiotic flies (80). These gnotobiotic flies were then placed on either a control (no insecticide) or toxic (insecticide-treated) diet, with or without microbiota transplants. This resulted in four experimental conditions: 1) control flies on control media, 2) control flies on toxic media, 3) resistant flies on control media, and 4) resistant flies on toxic media. While axenic flies were also reared on control media as part of the experimental design, these groups were not central to our primary analysis and are not presented in the main text (see Dryad repository for full datasets). Microbiota transplants were used to test whether microbiome evolution contributed to host adaptation. Gnotobiotic flies in each condition received one of two gut microbiota sources: 1) a pooled homogenate from indoor-reared flies originating from control populations or 2) a pooled homogenate from indoor-reared flies originating from resistant populations. These donor populations were derived from the outdoor experimental mesocosms but were reared in a common garden for two generations before microbiota extractions to ensure that any observed effects reflected evolved microbial communities rather than transient environmental effects.

Gut tracts from 20 adult flies in respective control or resistant donor populations were dissected, pooled, and homogenized in 700 μL of autoclaved 5% sucrose solution on ice (81), followed by bead-beating for approximately 30 seconds. Gnotobiotic eggs were inoculated with 50 μL of the homogenate corresponding to their assigned microbiota source (45). Separate pools of flies used for homogenate preparation were extracted for 16S sequencing to confirm the microbiota composition of donor populations.

Flies were maintained at 25°C on a 14L:10D cycle throughout the experiment. Egg-to-adult survivorship was recorded at 12-h intervals post-oviposition, with a final count performed after freezing all samples at −80°C. Body mass was measured by collecting pools of five females from each treatment, drying them at 55°C for 24 hours, and calculating the average individual mass. DNA was extracted from males pooled across replicates to confirm successful microbiota transplant and gnotobiotic rearing.

### Quantification of microbial communities from experimental treatments

To analyze the gut microbial communities of *D. melanogaster* at the conclusion of the experiment, DNA extractions were performed using the NucleoSpin Soil DNA kit (Macherey-Nagel, Düren, Germany). Each gut was pressed to begin tissue breakup prior to bead beating according to kit instructions. DNA was eluted using 50 µL of 5 mM Tris/HCl at pH 8.5. Subsequent DNA quantification was conducted with 5 µL per sample of eluted DNA on a Qubit (Invitrogen, Massachusetts, United States) using a dsDNA high sensitivity assay. With every set of extractions completed, blank samples were additionally run for downstream analysis of the kit’s sterility. Samples that had low DNA concentrations were vacufuged before sequencing to increase concentration. Isolated DNA was quantified (Equalbit 1x dsDNA HS Assay kit) and amplified using primers covering the V3-V4 hypervariable 16s rRNA region. Library quality was assessed and libraries were dual indexed before Equimolar pooling based on QC values. Pooled libraries were sequenced on an Illumina MiSeq with a 250 bp read length configuration to a depth of 0.3M reads for each sample.

### 16S rRNA sequencing and data processing

Data from the outdoor mesocosm experiment were imported as demultiplexed single-end reads with primers trimmed during importation. High-resolution denoising and filtering were performed using the DADA2 pipeline (82), with sequences truncated at a quality score of 20. Default QIIME2 parameters (83) were applied for filtering, yielding an initial dataset with an average input read count of 108,506 per sample. Post-filtering and chimeral-read removal retained a mean of 32,037 high-quality reads per sample, further reduced to 3,376 reads after excluding *Wolbachia* sequences, which were removed as they are intracellular (48, 84).

Data from the manipulative indoor experiment was imported using demultiplexed paired-end reads with primers trimmed during importation. High-resolution denoising and filtering were performed using the DADA2 pipeline (82), with sequences truncated at a quality score of 20. Default QIIME2 parameters (83) were applied for filtering, retaining 7.59% of input-reads after filtering. For consistency, *Wolbachia* sequences were also removed as they are intercellular (48, 84).

Taxonomic classification was performed using the Greengenes full-length 16S rRNA database with a scikit-learn naïve Bayes classifier (85). Retained reads primarily consisted of sequences from the families *Acetobacteraceae* and *Lactobacillaceae*, consistent with prior studies of *D. melanogaster* microbiomes (46, 48, 84). Aggregated taxonomic data at the family level were used to evaluate relative abundance and differential abundance patterns.

An unrooted phylogenetic tree was constructed to enhance the accuracy of inferences via weighted UniFrac analyses. Further filtering in *phyloseq* (86) excluded taxa with fewer than five total observations across all samples. Data from technical replicates (i.e., same cage, treatment, and exposure) were combined to reduce sampling error and improve statistical power. No rarefaction was performed. Beta diversity metrics (weighted UniFrac and Bray-Curtis) and alpha diversity indices were calculated to assess compositional and diversity shifts in microbiomes associated with insecticide exposure.

### Effect of microbiota transplant on recipient microbiomes

Transplant samples were compared to axenic samples to confirm that microbiota inoculation altered the microbial composition of recipient flies. No significant differences were observed on the ASV level (*t*_38,39_ = 0.281, P = 0.78). However, microbial read counts differed significantly between treatments (*t*_38,39_ = 6.46, P < 0.01), indicating that species diversity (AVG Transplant = 20, AVG Axenic = 18) did not differ substantially, but overall microbial presence was higher in transplants (AVG Reads = 6,169) than in axenic samples (AVG Reads = 376). Relative abundance analyses revealed compositional differences between transplant and founder microbiomes, driven by specific taxa. Within control exposures, transplants also exhibited significant shifts in microbial composition relative to founders. Microbial composition in gnotobiotic samples was significantly distinct from blank controls (F_1,23_ = 76.91, P < 0.01). Gnotobiotic samples had an average of 376.65 reads, while blank samples contained 6,707.4 reads, confirming successful sterilization and rearing of gnotobiotic flies on axenic media.

### Data analysis

All statistical analyses and plotting were conducted in R version 4.4.2 (87).

### Phenotypic evolution

To test for the effects of insecticide exposure on fitness-related traits, we used linear mixed-effects models executed in the *lme4* package (88). Each phenotype (survivorship, development rate, fecundity, starvation tolerance, and adult weight) was treated as an independent response variable, with ‘Time’ (*T*_0_ → *T*_4_), ‘Population’ (control vs. resistant), and their interaction as fixed effects. ‘Cage’ was included as a random effect to account for repeated measures within experimental replicates.

Significance was assessed using Type III ANOVA with Satterthwaite’s method for estimating denominator degrees of freedom, implemented via the *lmertest* package (89). This allowed us to test whether phenotypic changes occurred over time and whether resistant and control populations diverged. Pairwise contrasts between control and resistant populations at each time point were performed using the *emmeans* package (90), with Benjamini-Hochberg correction applied to adjust for multiple comparisons (91).

### Microbiome evolution

To assess whether insecticide exposure drove microbiome evolution, we tested for differences in survival, development rate, and adult weight between microbiomes from control and resistant populations in axenic and xenic conditions. For each phenotype, we first evaluated the normality of the model residuals using the Shapiro-Wilk test. Based on these results, we applied either a one-sample *t*-test to determine whether the mean difference between microbiome treatments deviated significantly from zero, or a Wilcoxon signed-rank test when normality assumptions were violated. These analyses allowed us to determine whether microbiome-associated changes in host phenotypes were consistent with evolutionary shifts rather than plastic responses.

### Microbiome transplant

To test whether microbiome evolution contributed to host phenotypic adaptation, we analyzed survival, development rate, and adult weight in flies receiving microbiota transplants from either control or resistant donor populations. We used linear models to test for effects of microbiome donor source and rearing environment on each phenotype. To evaluate overall differences, we conducted ANOVA on model outputs. Post-hoc pairwise comparisons were performed using Tukey’s HSD test to assess specific contrasts between microbiome donor treatments. This approach allowed us to test whether microbiome transplants influenced host phenotypes independently of plastic responses to insecticide exposure.

### Evolutionary rate calculation

We calculated evolutionary rates in Haldanes to quantify the rate of phenotypic change over the experiment. We calculated the mean Haldane’s rate across replicate cages within each population to obtain a single rate per time interval. Evolutionary rates were calculated for each experimental interval (*T*_0_ → *T*_1_, *T*_1_ → *T*_2_, *T*_2_ → *T*_3_, *T*_3_ → *T*_4_) and for the entire experiment (*T*_0_ → *T*_4_).

### Microbiome sequences

Microbiome sequencing data were analyzed using the *phyloseq* (86) and *vegan* (92) packages in R. Alpha diversity (Shannon diversity index) was calculated for each sample, and the effects of treatment, media type, and their interaction were tested using LMEs. Beta diversity metrics, including Bray-Curtis dissimilarity and weighted UniFrac distances, were used to evaluate microbial community structure. Permutational multivariate analysis of variance (PERMANOVA) was performed using the *adonis2* function in *vegan*, with data subset to independent ‘population’ and ‘treatment’ as fixed effects.

To analyze bacteria abundance across treatments, the most abundant family-level taxa (*Acetobacteraceae* and *Lactobacillaceae*) were isolated and analyzed independently across treatments using a generalized linear models (GLMs) with a Poisson family to identify taxa showing differential abundance across groups.

To determine whether microbiomes changed in parallel across independent populations, we quantified parallelism using the *multivarvector* package (93). Multivariate vectors connecting mean Principal Coordinate Analysis (PCoA) scores for control and insecticide-exposed samples were calculated to estimate the magnitude and direction of microbiome shifts in response to insecticide exposure. Angular values between vectors were used to assess parallel, non-parallel, or anti-parallel relationships among replicate cages.

## Data accessibility statement

The data supporting the results are archived in the public repository Dryad under DOI (insert DOI link here). This link has been created to be shared during the peer-review process to allow access only to the data files included with our submission.

## Competing interest statement

The authors declare no competing interests.

## Acknowledgements

We thank S. Porter and the Rudman lab for helpful comments, and B. Campagnari and C. Clare for assistance with some of the data collection. Participation in the 2023 Animal-Microbe Symbioses Conference hosted by the Gordon Research Conference was influential in the conceptual design of this research.

## Funding

This research was supported by the National Institute of General Medical Sciences of the National Institutes of Health under Award Number R35GM147264 (S.M.R.). The content is solely the responsibility of the authors and does not necessarily represent the official views of the National Institutes of Health. Additional support was provided by a Natural Sciences and Engineering Research Council of Canada (NSERC) Postdoctoral Fellowship and a Liber Ero Postdoctoral Fellowship (R.S.S.).

## Notes

### Competing Interest Statement

The authors have declared no competing interest.

