## Supporting Information for "Microbiome evolution plays a secondary role in host rapid adaptation"

**This file includes:** Figure S1-S3, Table S1, Table S2

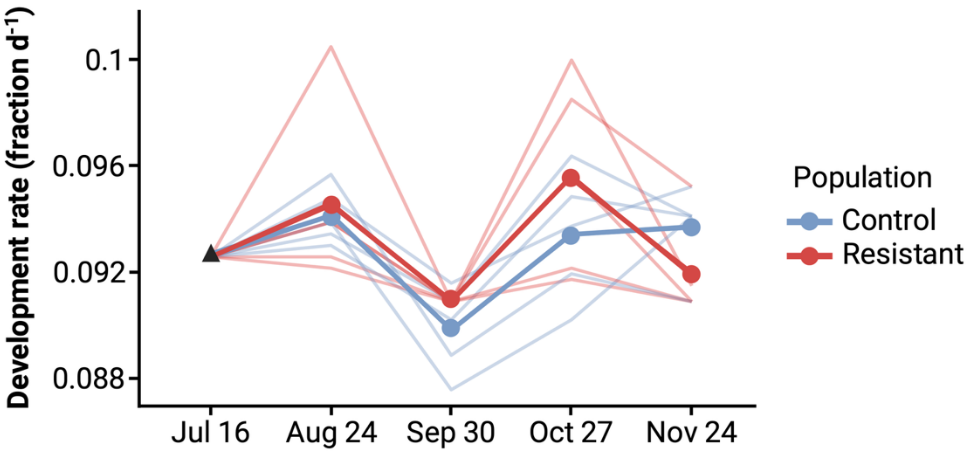


**Figure S1.** Phenotypic evolution of development rate in populations exposed (‘resistant’) and unexposed (‘control’) to the insecticide. Development rate trajectories were measured at each time point following two generations of common garden rearing, with all assays conducted on insects reared on toxic (insecticide) media to assess resistance. Mean development rate was calculated as the fraction of development time completed per day [1/(total hours/24)]. The black triangle (▲) represents the mean development rate of the founding population (outdoor cages initiated July 16), measured under toxic assay conditions. Thin colored lines represent the mean development rate trajectory of each individual population, and thick colored lines show the treatment means (averaged across cages).

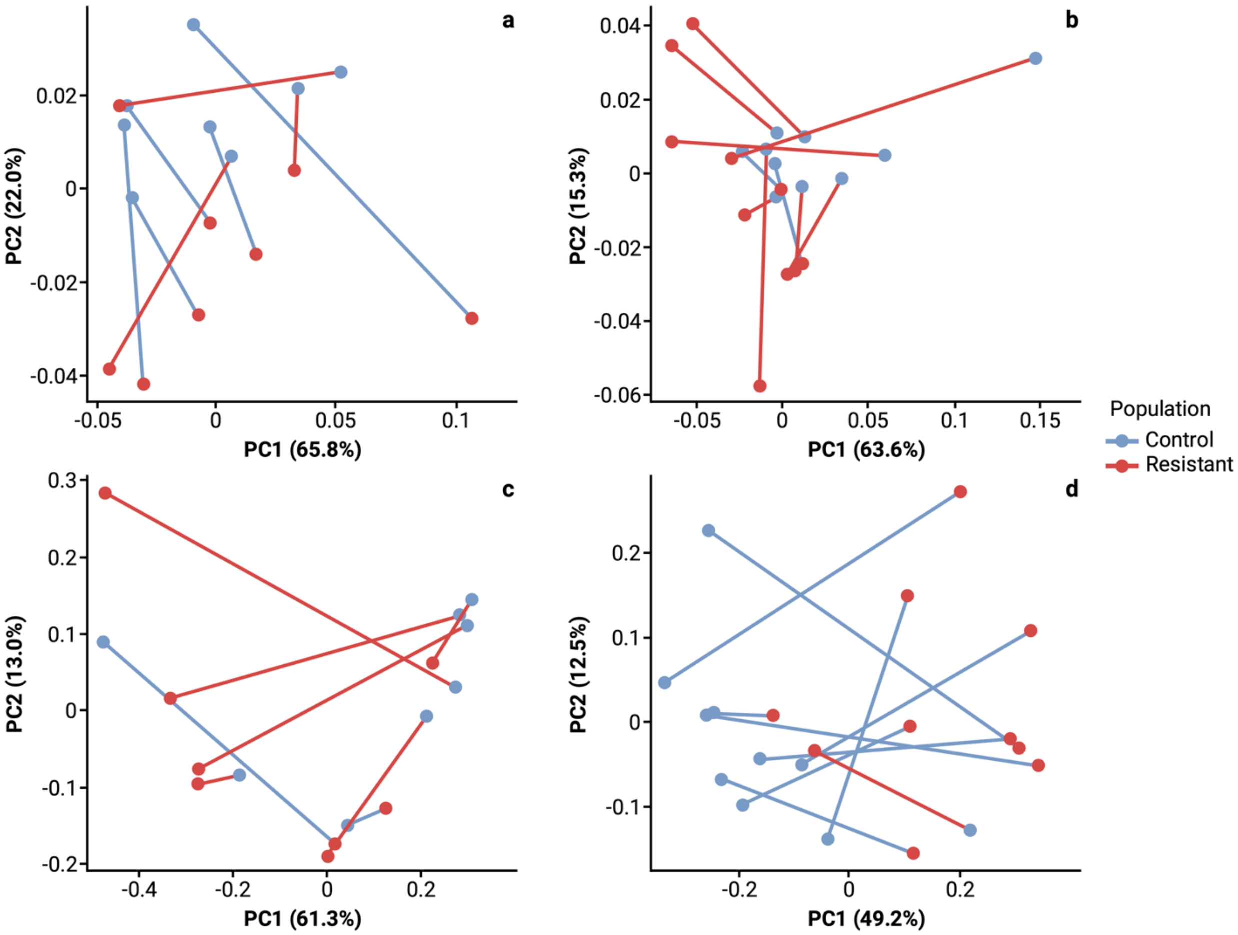


**Figure S2.** Test for parallelism in the shifts of microbiome composition during resistance evolution. Vectors represent compositional changes in individual replicate populations between founder and October 27, following resistance evolution, plotted using principal coordinate analysis (PCoA) based on Weighted UniFrac (A-B) and Bray-Curtis (C-D) distance matrices. Each panel shows replicate trajectories for resistant and control populations. Angles between replicate trajectories were used to quantify parallelism (reported in the main text).

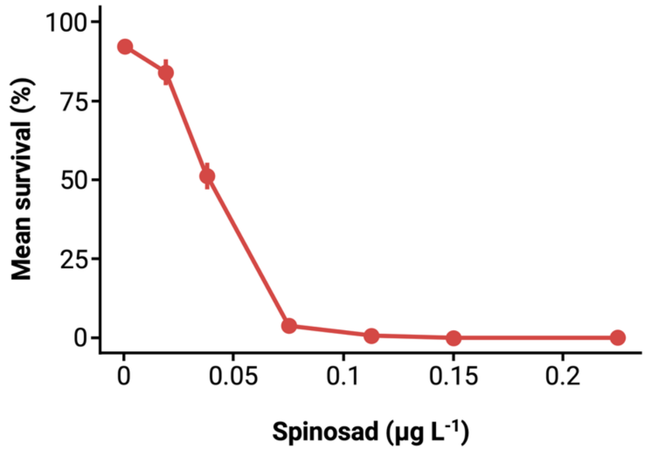


**Figure S3.** Dose-response relationship for egg-to-adult survivorship in *Drosophila melanogaster* exposed to the insecticide spinosad. Mean survival (± 1 SE) across a range of spinosad concentrations (µg $L^{-1}$) are plotted, and the lethal dose at which 50% of *D. melanogaster* experienced mortality was estimated and used to inform the experimental concentrations used in the present work (see Methods).

**Table S1.** Mean evolutionary rates (Haldanes) for life-history traits across timepoints. Mean haldane values (standardized evolutionary rates per generation) calculated from $T_0 \rightarrow T_1$, $T_1 \rightarrow T_2$, $T_2 \rightarrow T_3$, $T_3 \rightarrow T_4$, and $T_0 \rightarrow T_4$ for survivorship, development rate, fecundity, starvation tolerance, and adult weight.

***Survival***

| Population | $T_0 \rightarrow T_1$ | $T_1 \rightarrow T_2$ | $T_2 \rightarrow T_3$ | $T_3 \rightarrow T_4$ | $T_0 \rightarrow T_4$ |
|---|---|---|---|---|---|
| Control | 0.3806 | -0.3486 | 0.356972 | -0.4038 | 0.022276 |
| Resistant | 0.8878 | -0.315 | 0.3956 | -0.2474 | 0.2848 |

***Development rate***

| Population | $T_0 \rightarrow T_1$ | $T_1 \rightarrow T_2$ | $T_2 \rightarrow T_3$ | $T_3 \rightarrow T_4$ | $T_0 \rightarrow T_4$ |
|---|---|---|---|---|---|
| Control | 0.694 | -1.05 | 0.583 | 0.0428 | 0.107 |
| Resistant | 0.281 | -0.516 | 0.602 | -0.424 | 0.0582 |

***Fecundity***

| Population | $T_0 \rightarrow T_1$ | $T_1 \rightarrow T_2$ | $T_2 \rightarrow T_3$ | $T_3 \rightarrow T_4$ | $T_0 \rightarrow T_4$ |
|---|---|---|---|---|---|
| Control | -0.07396 | 0.3392 | -0.107258 | -0.07538 | 0.013566 |
| Resistant | 0.0531256 | 0.20972 | -0.08884 | 0.12966 | 0.07032 |

***Starvation tolerance***

| Population | $T_0 \rightarrow T_1$ | $T_1 \rightarrow T_2$ | $T_2 \rightarrow T_3$ | $T_3 \rightarrow T_4$ | $T_0 \rightarrow T_4$ |
|---|---|---|---|---|---|
| Control | -0.1553 | -0.19066 | 0.00346 | 0.11634 | 0.08382 |
| Resistant | -0.04972 | -0.2132 | 0.05238 | 0.3898 | 0.044 |

***Adult weight***

| Population | $T_0 \rightarrow T_1$ | $T_1 \rightarrow T_2$ | $T_2 \rightarrow T_3$ | $T_3 \rightarrow T_4$ | $T_0 \rightarrow T_4$ |
|---|---|---|---|---|---|
| Control | 1.2975 | -0.5783333 | -0.739175 | -0.1288 | 0.08938 |
| Resistant | 1.4316 | -0.16664 | -0.3162 | -0.1334 | 0.18204 |

**Table S2.** Relative abundance of bacterial families detected in *Drosophila melanogaster* microbiomes. Complete list of microbial families detected in control and resistant populations following two generations of common garden rearing, used for analyses of microbiome response to insecticide exposure and evolution of resistance.

| **Family** | **Read count (ASV)** |
|---|---|
| Acetobacteraceae | 545,670 |
| Lactobacillaceae | 4,839 |
| Staphylococcaceae | 303 |
| Sphingomonadaceae | 216 |
| Rhodobacteraceae | 157 |
| Micrococcaceae | 108 |
| Marinilabiliaceae | 91 |
| Clostridiaceae | 66 |
| Mycobacteriaceae | 48 |
| Neisseriaceae | 46 |
| Chroococcidiopsidaceae | 39 |
| Moraxellaceae | 38 |
| Blastocatellaceae | 19 |
| Streptococcaceae | 16 |
| Caulobacteraceae | 15 |
| Deinococcaceae | 14 |
| Atopobiaceae | 14 |
| Beijerinckiaceae | 12 |
| Planococcaceae | 12 |
